# Alfalfa vein mottling virus, a novel potyvirid infecting *Medicago sativa* L

**DOI:** 10.1101/2023.09.07.556681

**Authors:** Lev G. Nemchinov, Olga A. Postnikova, William M. Wintermantel, John C. Palumbo, Sam Grinstead

## Abstract

We have recently identified a novel virus detected in alfalfa seed material. The virus was tentatively named alfalfa-associated potyvirus 1, as its genomic fragments bore similarities with potyvirids. In this study, we obtained a complete genome sequence of the virus and classified it as a tentative species in the new genus, most closely related to the members of the genus *Ipomovirus* in the family *Potyviridae*. This assumption is based on the genome structure, phylogenetic relationships, transmission electron microscopy investigations and, in part, on serological cross-reactivity of the virus. We also demonstrated its mechanical transmission to the indicator plant *Nicotiana benthamiana* and to the natural host *Medicago sativa*, both of which developed characteristic symptoms therefore suggesting a pathogenic nature of the disease. Consistent with symptomatology, the virus was renamed to alfalfa vein mottling virus. A name Alvemovirus was proposed for the new genus in the family *Potyviridae*, of which alfalfa vein mottling virus is a tentative member.

## Introduction

Research on alfalfa (*Medicago sativa* L.), a major forage crop worldwide, greatly benefited from the latest advances in high throughput sequencing (HTS) technologies that enabled the completion of its genome assembly [1] revealing new insights into the mechanisms of the crop’s resistance to abiotic and biotic stresses [2,3,4,5], identification of genetic markers associated with important agronomic traits [6,7,8] and discovery of numerous microorganisms infecting alfalfa, primarily viruses [9–12]. Viruses were found to be an integral part of multi-pathogenic infections collectively forming the alfalfa pathobiome, a diverse community of pathogenic microbes within the biotic environment of the plant [13,14]. Recently, during the initial seed screenings of alfalfa germplasm accessions maintained by the USDA ARS National Plant Germplasm System (NPGS), we have detected a broad range of viruses in the mature seeds of different germplasm sources [15]. Among them were fragments of a poty-like virus with a genomic organization typical for *Potyviridae* that shared ∼ 26%-32% identity with a few members of the family. Therefore, the virus was provisionally named alfalfa-associated potyvirus 1 (AaPV1), [15]. In the present study, aimed at surveying alfalfa varieties grown in the U.S. to identify, characterize, and prevent the spread of novel and emerging viruses in the country, AaPV1 was found in plant samples collected from commercial alfalfa fields in Arizona, USA. A coding-complete genome of the virus was assembled from overlapping sets of contigs and a phylogenetic relationships with other members of the family *Potyviridae* were established. The virus was further categorized as a novel species, tentatively related to the genus *Ipomovirus* and potentially representing a new genus in the family *Potyviridae*. Consistent with symptomatology in the natural host *Medicago sativa* L., the virus was designated as **al**falfa **ve**in **mo**ttling virus (AVMV), a representative member of a new taxon, provisionally named Alvemovirus.

## Materials and Methods

### Plant material

Five alfalfa plants (stems and leaves) were sampled from each of the four different fields, 10-15 acres in size, located in Yuma Country, Arizona, USA. Geographic coordinates of the alfalfa fields and the adjacent crops are shown in Table 1.

**Table 1.** Geographic locations of alfalfa fields.

| Alfalfa fields, № | GPS coordinates |  | Adjacent crops | № of plants collected |
| --- | --- | --- | --- | --- |
| 24E | 32° 40' 56.58"N | 114° 13' 42.72"W | Cantaloupe, lettuce, cauliflower | 5 |
| 32E | 32° 42' 38.62"N | 114° 05' 28.21" W | Cantaloupe, cauliflower, spinach | 5 |
| 43E | 32° 47' 11.44"N | 113° 54' 15.81"W | Cantaloupe, lettuce, broccoli, cauliflower | 5 |
| 51E | 32° 46' 42.26"N | 113° 45' 20.89"W | Cantaloupe, lettuce | 5 |

### Total RNA extraction, RNA sequencing, RT-PCR, and 5’RACE

Five leaves were pooled from each of the 20 plants for RNA extraction. Total RNA was extracted using Promega Maxwell® RSC Plant RNA Kit (Promega Corp., Fitchburg, WI) according to the manufacturer’s directions. cDNA libraries were prepared using Illumina TruSeq Stranded Total RNA with the Ribo-Zero Library Prep Kit (Illumina Inc., San Diego, CA USA) and RNA sequencing was performed by Psomagen (Psomagen Inc., Rockville MD USA) on a NovaSeq6000 S4 platform (150PE, 1Gb, 20 million total reads). Reverse transcription–polymerase chain reactions (RT-PCR) were performed using the SuperScript One-Step RT-PCR System according to the manufacturer’s directions (Thermo Fisher Scientific Inc., Waltham, MA USA) or utilizing SuperScript™ III First-Strand Synthesis System Thermo Fisher Scientific, Waltham, MA USA) following by PCR amplification with AmpliTaq Gold DNA Polymerase Thermo Fisher Scientific). Total RNA was extracted using TRIzol Reagent (Thermo Fisher Scientific Inc., Waltham, MA USA). Negative control reactions were performed with RNA obtained from uninfected *N. benthamiana* or with sterile RNAse-free water processed for cDNA preparation the same way as RNA extracted from the infected alfalfa plants. Three virus-specific sets of primers were designed based on the results of the HTS analysis: 1) LN1077/LN1078 (forward, 5’ CTCCCTTTGGTACTCGGTATTG, 3′, pos. 7679-7701 and reverse, 5′ CTCTAATGTGCGGACCTTTCT 3′, pos. 8072-8093, size 414 bp); 2) LN1079/LN1080 (forward, 5′ TGACAGCGAGTTCTCATTCC 3′, pos. 9165-9184, and reverse, 5′ GTCAGTCAAACCAGCCTTTATTC 3′, pos. 9800-9822, size 658 bp); 3) LN1095/LN1096 (forward, 5′ CCAGGGTGTTGTTGATGATTTG 3′, pos. 4176-4197, and reverse, 5′ TCTCAGTCACATTCCGCATAAA 3′, pos. 4579-4600, size 425 bp). PCR products were either sequenced directly or cloned into the TOPO II vector (Thermo Fisher Scientific Inc., Waltham, MA USA) and sequenced. 5’RACE was performed with SMARTer RACE 5’/3’ Kit (Takara Bio USA, Inc. San Jose, CA USA) according to the manufacturer’s directions.

### Bioinformatic analysis

A schematic representation of the bioinformatics pipeline used in this study is shown in **Fig.1**. Briefly, sequence reads were first trimmed using Trimmomatic [16], followed by their assembly with SPAdes [17]. The resulting contigs were screened using BLASTx searches [18] against a virus database containing all plant virus protein sequences from the NCBI RefSeq database (https://www.ncbi.nlm.nih.gov/refseq/). The potential plant viral hits were searched once again using BLASTx against the full NCBI nr protein database. BBMap [19] was used to generate sequencing coverage values for the identified contigs. For the prediction of cleavage sites for viral proteases, polyproteins from 109 different members of the family *Potyviridae* were aligned by four different Multiple Sequence Alignments (MSA) software applications: ClustalW, ClustalO, MUSCLE, and CLC Genomics Workbench 23, very accurate setting (QIAGEN LLC, Germantown MD USA). Cleavage sites obtained by each MSA method were compared and the most likely sites were selected by another comparison with previously reported consensus sites [20].

**Figure 1.**
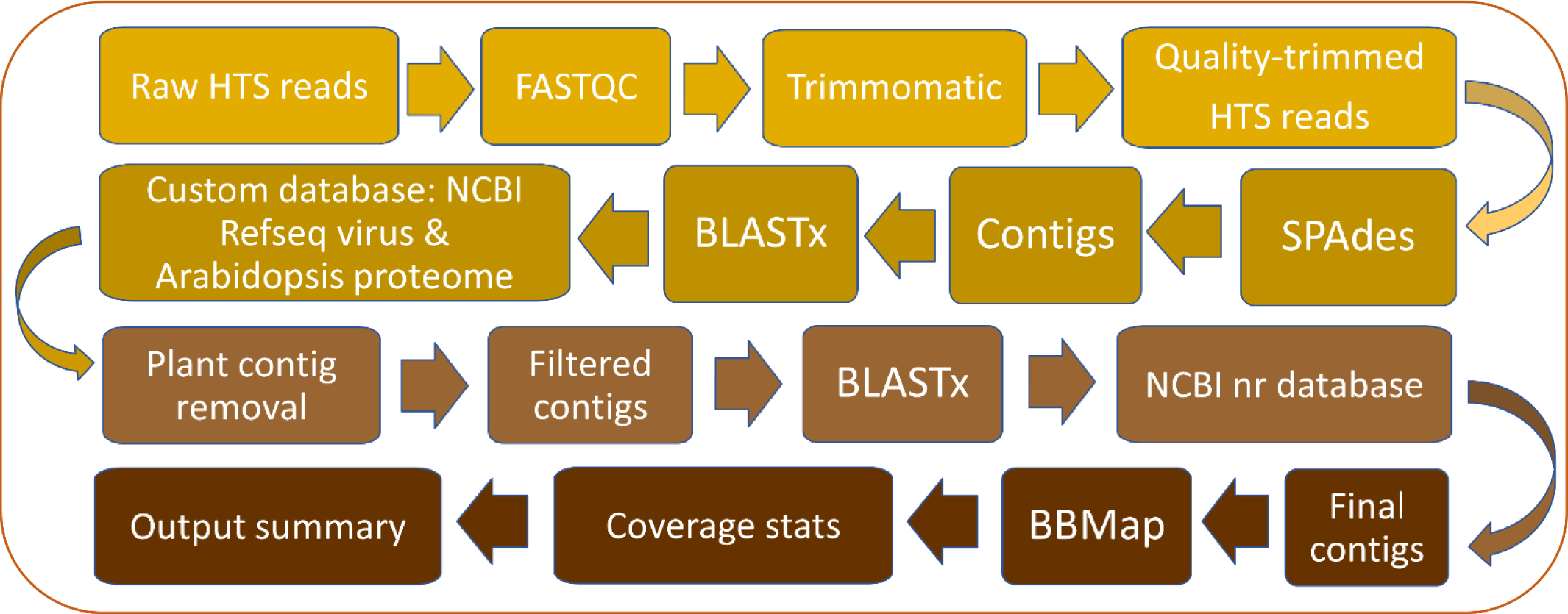
Bioinformatics pipeline

### Phylogenetic analysis

Phylogenetic analysis was performed with polyproteins from selected members of the family *Potyviridae*. Sequences were aligned with Clustal W and the unrooted tree was built with RAxML-NG [21] using the maximum likelihood algorithm, a maximum of 1000 bootstrap replicates, and bootstopping (autoMRE, cutoff: 0.030000). Boostrapping converged after 150 replicates.

### Mechanical inoculation of *Nicotiana benthamiana* Domin. and alfalfa (*Medicago sativa* L.) plants

Symptomatic leaves from field alfalfa plants were homogenized in cold 20 mM potassium phosphate buffer, pH 7.0, using a sterile mortar and pestle. The resulting suspension was briefly centrifuged to precipitate plant debris and ten microliters of the supernatant were rubbed onto carborundum-dusted leaves of three-weeks old *N. benthamiana* plants using sterile inoculating loops. Carborundum-dusted leaves of the three weeks-old seedlings of two alfalfa cultivars, cv. CUF 101 and cv. Maverick, were rub-inoculated with purified viral preparations using sterile inoculating loops.

### Virus purification

The virus was partially purified from 20 grams of symptomatic *N. bentamiana* leaves using a protocol developed for *Poinsettia mosaic virus* [22].

### Transmission electron microscopy

For transmission electron microscopy (TEM), TEM grids were incubated on the drop of purified viral preparation for 2 minutes, rinsed with one milliliter of sterile water and stained with 1% phosphotungstic acid (PTA) solution. The grids were examined in a Hitachi H-7700 Electron Microscope at the Electron and Confocal Microscope Unit, Beltsville Agricultural Research Center.

### Western blot assays

Western blots (WB) were performed as previously described [23]. Membranes were probed with polyclonal antiserum to sweet potato mild mottle virus (SPMMV; DSMZ, Braunschweig, Germany) using the company’s recommended dilution (1:1000).

## Results

### Symptoms displayed on the infected alfalfa plants

According to the results of HTS, eight out of 20 plants examined (40%) originating from three different alfalfa fields, were infected with the virus resembling AaPV1. These plants displayed a variety of virus-like symptoms including mottling, chlorosis, vein clearing, and bright yellow blotches (**Fig. 2**). The symptoms were likely triggered by multiple co-infecting pathogens, including the virus in question [14].

**Figure 2.**
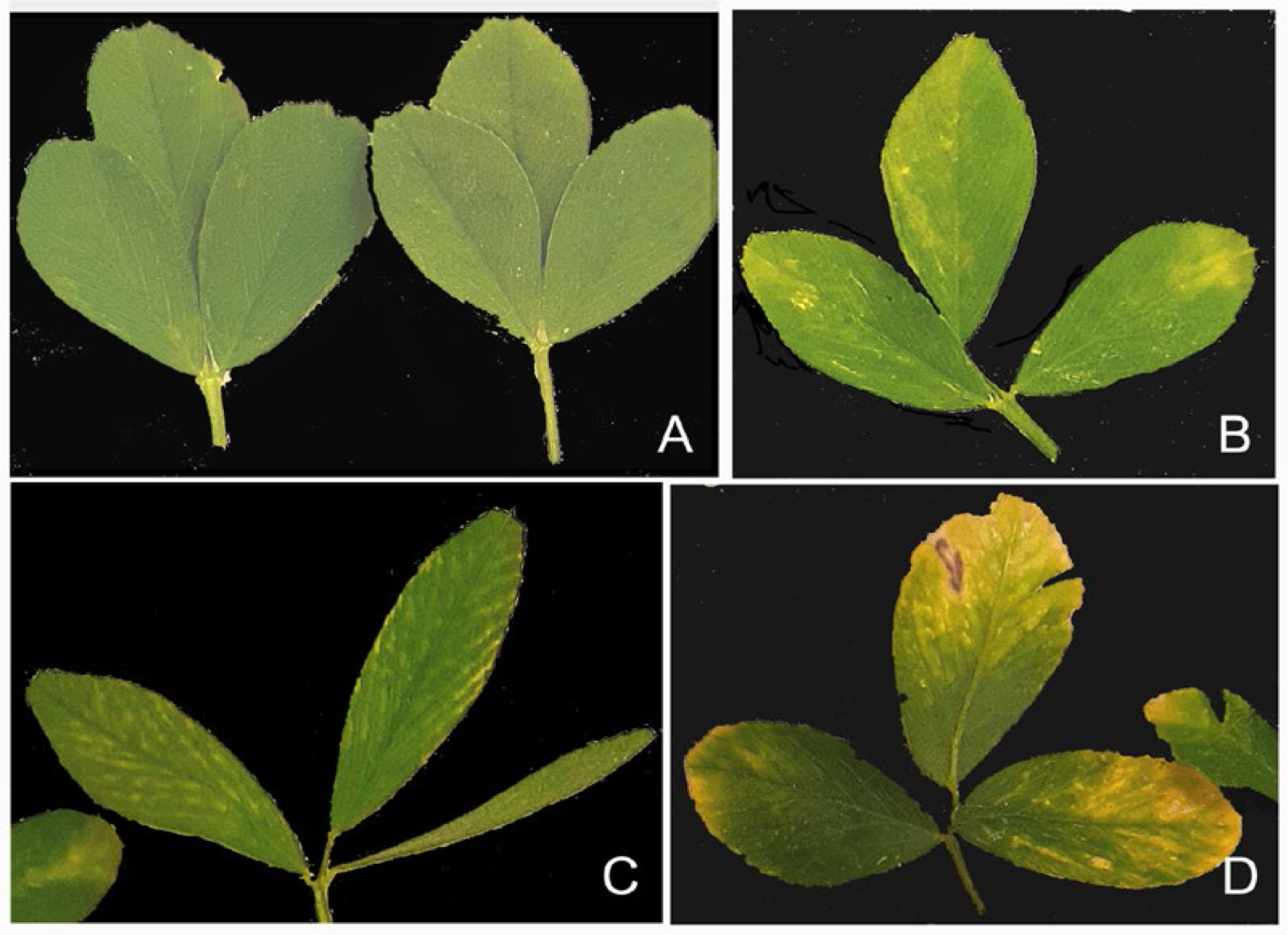
Diverse symptomatology of alfalfa plants in the field. **A**, asymptomatic plants. **B-D**, plants displaying different virus-like symptoms.

### Viral genome assembly and structure

Assembled contigs generated from each of the eight plants exceeded 9-10 kb in length and represented variants of the same virus (**Table S1**). The longest contig with 4940 x per-nt average coverage consisted of 10,323 nucleotides (nt) and included a poly (A) tail, thus indicating that the 3’-terminal portion of the viral genome is complete. Multiple 5’RACE reactions demonstrated that the HTS-derived length of the viral genome is 10,037 nt, excluding the poly(A) tail (75 nt). According to the sequenced 5’RACE PCR products, the 5’ non-coding region of the virus contains 107 nt; an open reading frame (ORF) encoding a large polyprotein comprises 9,732 nt (108-9,839 nt); and the 3’ non-coding region of the virus appears complete, encompassing 198 nt (9,840-10,037) without the poly (A) tail.

In a PSI-BLAST search, the top three hits for the viral polyprotein (excluding AaPV1 submissions) were: eggplant ipomovirus A (GenBank ID: BCG55390.1; 32.8% identity, query cover 78%; E-value=0.0); the roymovirus, rose yellow mosaic virus (GenBank ID: BBI90117.2; 31.6% identity; query cover 87%; E-value=0.0); and *Potyvirus* spp. polyprotein (GenBank ID: QHB15167.1; 31.3% identity, query cover 85%; E-value=0.0). Among other hits were squash vein yellowing virus (genus *Ipomovirus*; GenBank ID: AEV45694.1; 34,4% identity; query cover 65%, E-value=0.0); chili ringspot virus (genus Potyvirus; GenBank ID: APW85806.1; 28.7% identity; query cover 79%; E-value=0.0), and numerous species from different genera of the family *Potyviridae*.

The InterPro tool for functional analysis of proteins [24], predicted nine protein domains in the viral ORF: Potyviridae P1 protease domain (IPR002540); Helper component proteinase domain (PR001456); Protein P3 of potyviral polyprotein (IPR039560); Superfamilies 1 and 2 helicase ATP-binding type-1 domain (IPR014001); Superfamilies 1 and 2 helicase C-terminal domain (IPR001650; Potyviridae polyprotein domain (IPR013648); Potyvirus NIa protease (NIa-pro) domain (IPR001730); Viral RNA-dependent RNA polymerase domain (IPR001205); and Potyvirus coat protein domain (IPR001592). The tentative genome structure of the virus and predicted cleavage sites, obtained by multiple alignment of 109 different potyvirids, are shown in **Fig. 3**.

**Figure 3.**
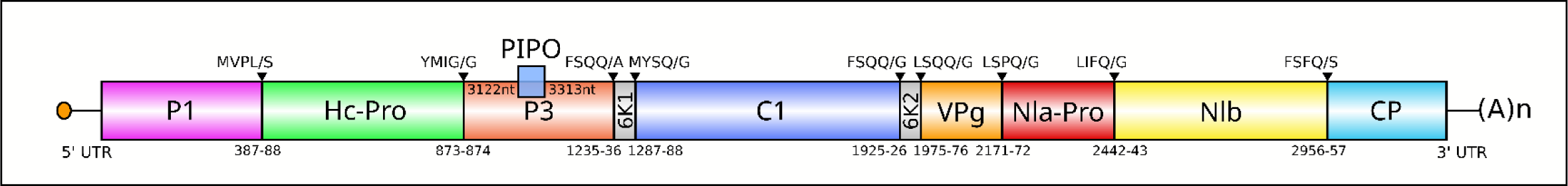
Tentative genome structure of the new virus infecting alfalfa. Black arrow heads indicate putative cleavage sites in the polyprotein. P1: protein 1 protease; P3: protein 3; PIPO: Pretty Interesting Potyviridae ORF (Chung et al. 2008); 6K1: 6 kDa peptide 1; CI: cylindrical inclusion; 6K2: 6 kDa peptide 2; VPg: virus protein genome-linked; Nia-Pro: nuclear inclusion a protease; NIb: nuclear inclusion b, RNA-directed RNA polymerase; CP: coat protein.

The virus does not appear to have a highly conserved G_1-2_ A_6-7_ motif preceding overlapping PIPO ORF characteristic for *Potyviridae* and located within the P3 cistron [25], Instead, there is an A_7_ motif, which is embedded in the overlapping ORF, presumably (according to the ORF Finder; https://www.ncbi.nlm.nih.gov/orffinder/) starting at the position 3122 nt (+3 ORF relative to the polyprotein) and followed by 63 codons with a termination codon TGA at pos. 3311-3313.

### Phylogenetic analysis

Phylogenetic analysis deduced from the alignment of the viral polyprotein and representative polyproteins of different members of the family *Potyviridae*, clustered the virus into a distinct outgroup, branching from the genus *Ipomovirus* (**Fig. 4**). This concurs with the significant divergence of the viral genomic and amino acid sequences from other members of the genus and likely indicates that the virus should be classified as member of a new, not yet established, genus in the family *Potyviridae*.

**Figure 4.**
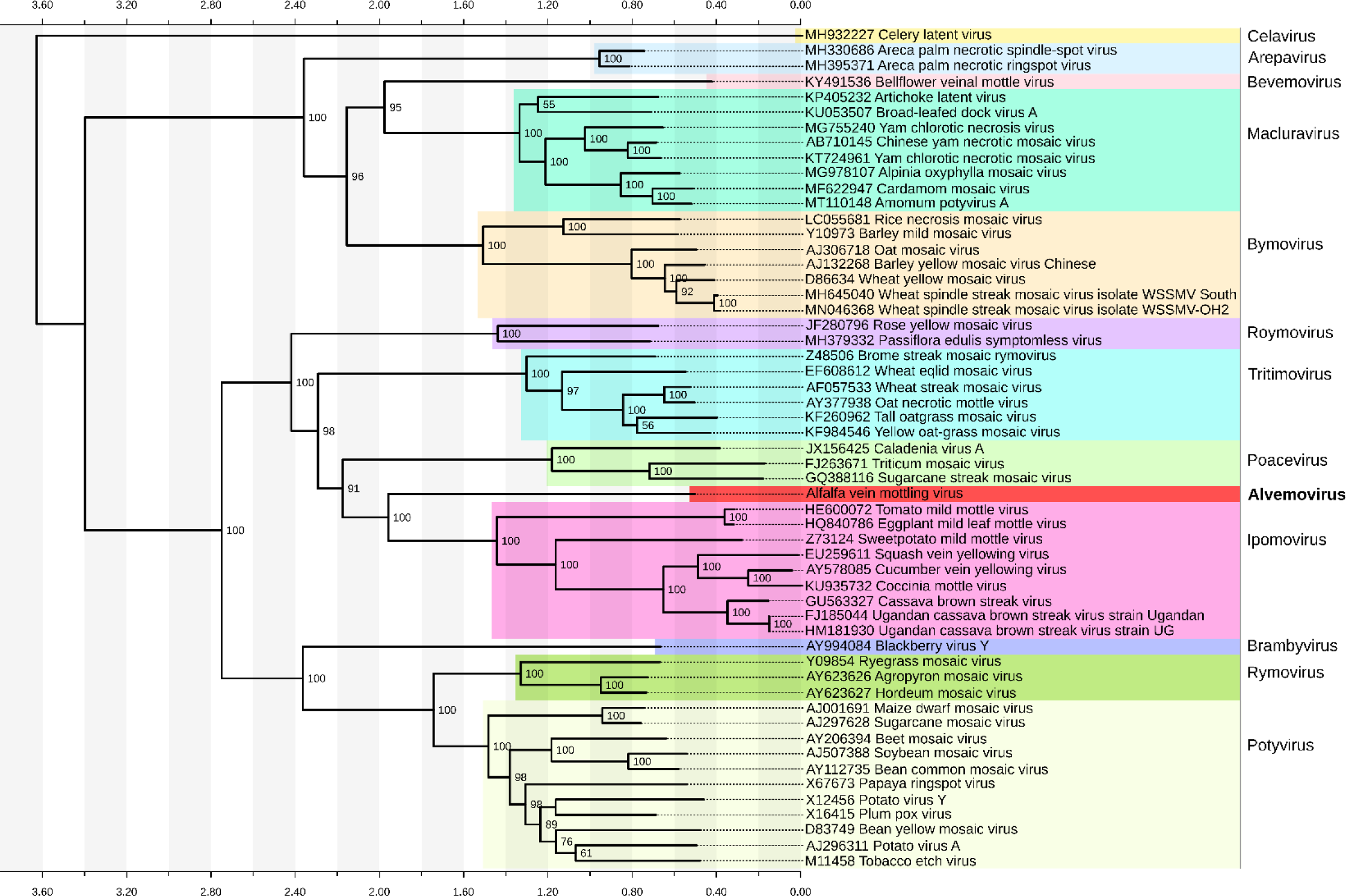
Phylogenetic relationships of alfalfa vein mottling virus with other members of the family *Potyviridae*. The unrooted tree was deduced from the Clustal W alignment and built with RaxML-NG [18] (Kozlov et al. 2019) tool using maximum likelihood algorithm, 1000 bootstrap replicates and bootstopping (autoMRE, cutoff: 0.030000).

### Mechanical transmission to *N. benthamiana* plants

Since potivirids are readily transmissible mechanically (https://ictv.global/report/chapter/potyviridae/potyviridae), we attempted mechanical transmission of the virus to *N. benthamiana*, a common indicator host for diagnosis of many plant viruses [26] (Hull, 2002). *N. benthamiana* plants mechanically inoculated with sap produced from infected alfalfa leaves, developed distinct symptoms two-three weeks post inoculation. The resulting symptoms could be visually characterized as chlorosis, mottling, and chlorotic lesions spots on non-inoculated leaves. The infected plants were also stunted compared to healthy control plants (**Fig. 5**). The infection of *N. bentamiana* plants was additionally confirmed by RT-PCR assay with primers LN1095/96 developed based on the HTS data and specific for the virus (**Fig.6**). The amplicon was of the correct size predicted by the analysis of the HTS-derived assembly and Sanger sequencing verified it is a fragment of the virus.

**Figure 5.**
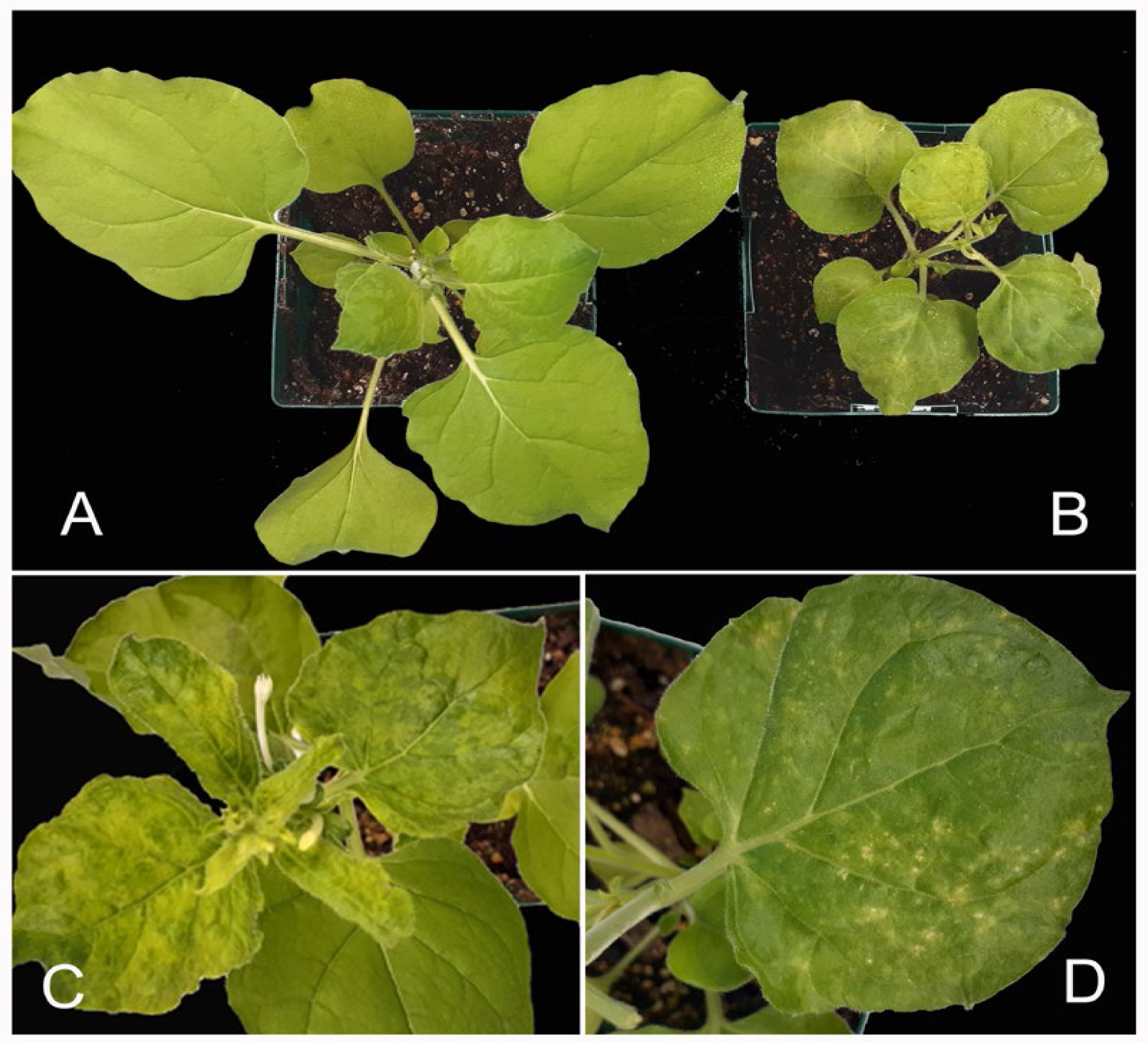
Symptoms developed on *Nicotiana benthamiana* plants after mechanical inoculation with AVMV-infected alfalfa. **A**, uninfected *N. benthamiana* plant. **B-D**, symptomatic plants.

**Figure 6.**
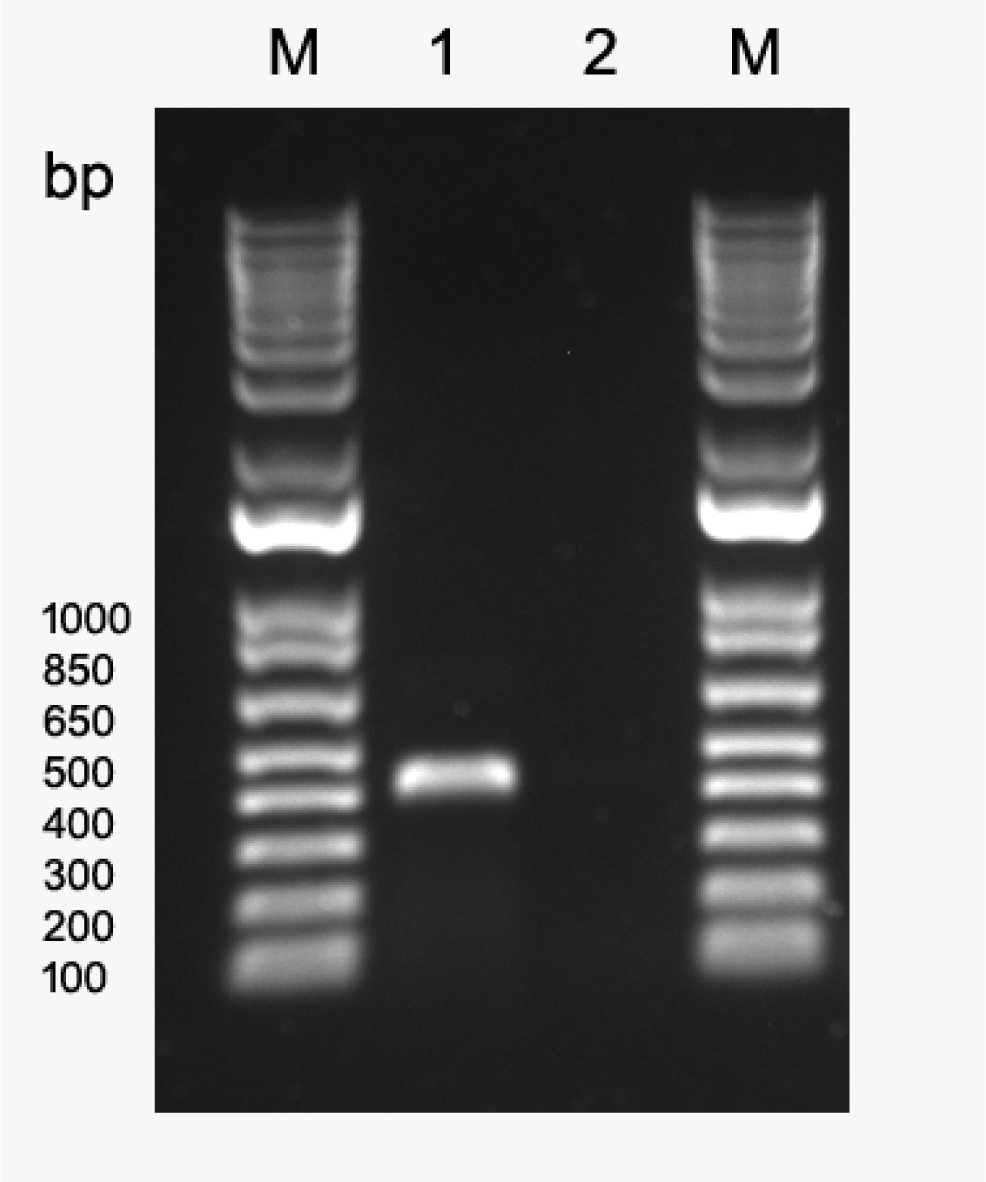
RT-PCR assay with total RNA extracted from symptomatic *Nicotiana benthamiana* plants. Virus-specific primers used in the assay were developed based on the HTS data. **M**, 1 Kb Plus DNA Ladder (Thermo Fisher Scientific, Waltham, MA USA). **1**, RT-PCR amplicon obtained from the virus-infected plant using primer pair LN1095/LN1096. **2**, no RT-PCR amplicons were obtained from the uninfected *N. benthamiana* plants.

### Transmission electron microscopy of the purified viral preparations

Transmission electron microscopy (TEM) can provide reliable information on particle dimensions and morphology, based on which a virus can be placed to a particular taxonomic group [27]. TEM examinations showed that the virions were flexuous filaments with a modal length of 800-1050 nm and the modal diameter of about 11–13 nm (**Fig.7**). This morphology is indicative of the members of the family *Potyviridae* and the larger particle length is consistent with the viruses of the genus *Ipomovirus*.

**Figure 7.**
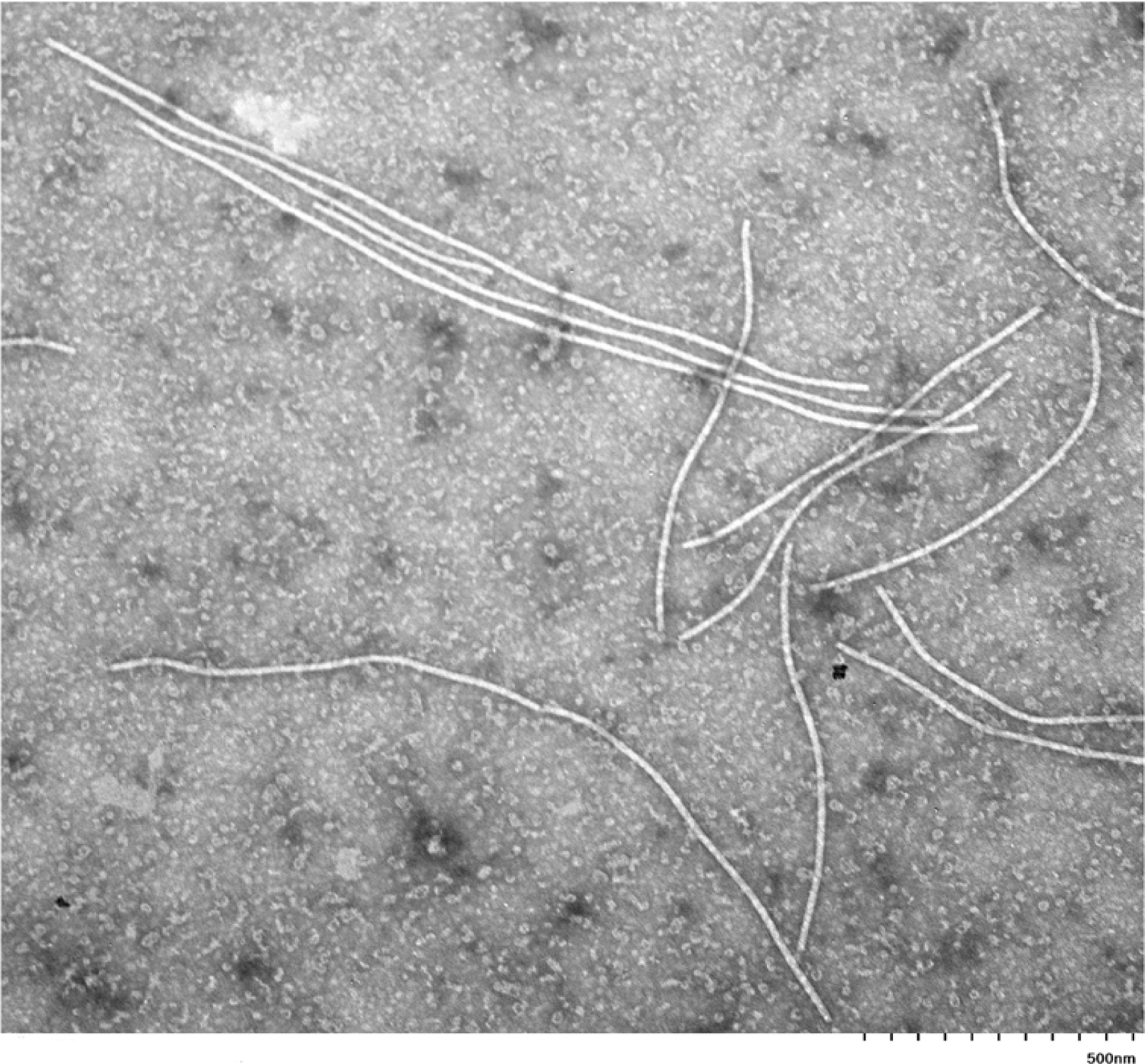
Transmission electron microscopy photograph of partially purified viral preparations. Magnification: 30,000. Scale bar: 500 nm.

### Mechanical transmission to alfalfa (*Medicago sativa* L.) plants

Among several plants of two inoculated alfalfa cultivars, only a single plant of cv. CUF 101 developed obvious symptoms ∼ 20 days post mechanical inoculation. Symptoms consisted of mottling and vein yellowing (**Fig. 8**). The symptomatic plant was tested, and the infection confirmed by RT-PCR assay with two different sets of primers developed based on the HTS data and specific for the virus (**Fig.9**). PCR products were sequenced and matched the virus in question.

**Figure 8.**
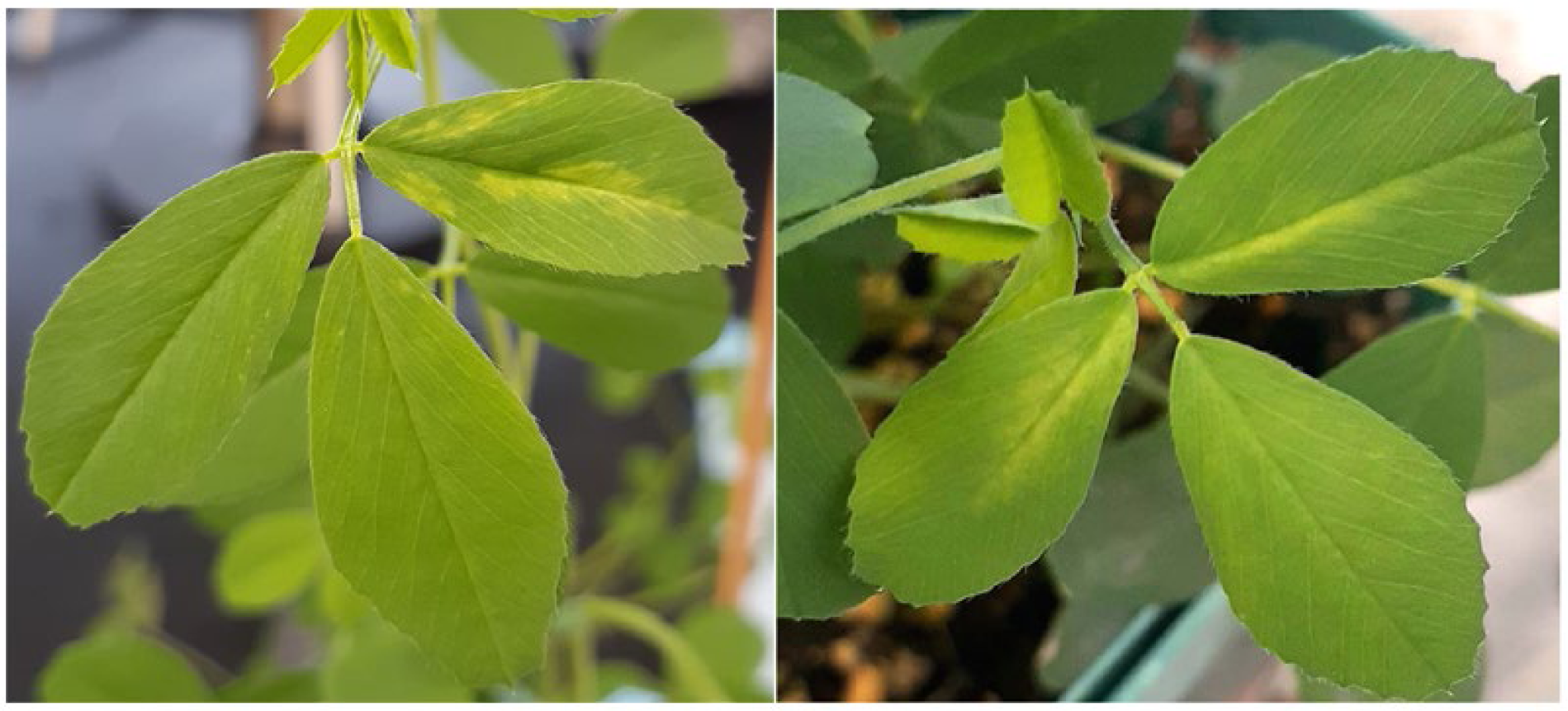
Symptoms developed on leaves of alfalfa (*Medicago sativa* L.) cv. CUF101 after mechanical inoculation with partially purified viral preparations.

**Figure 9.**
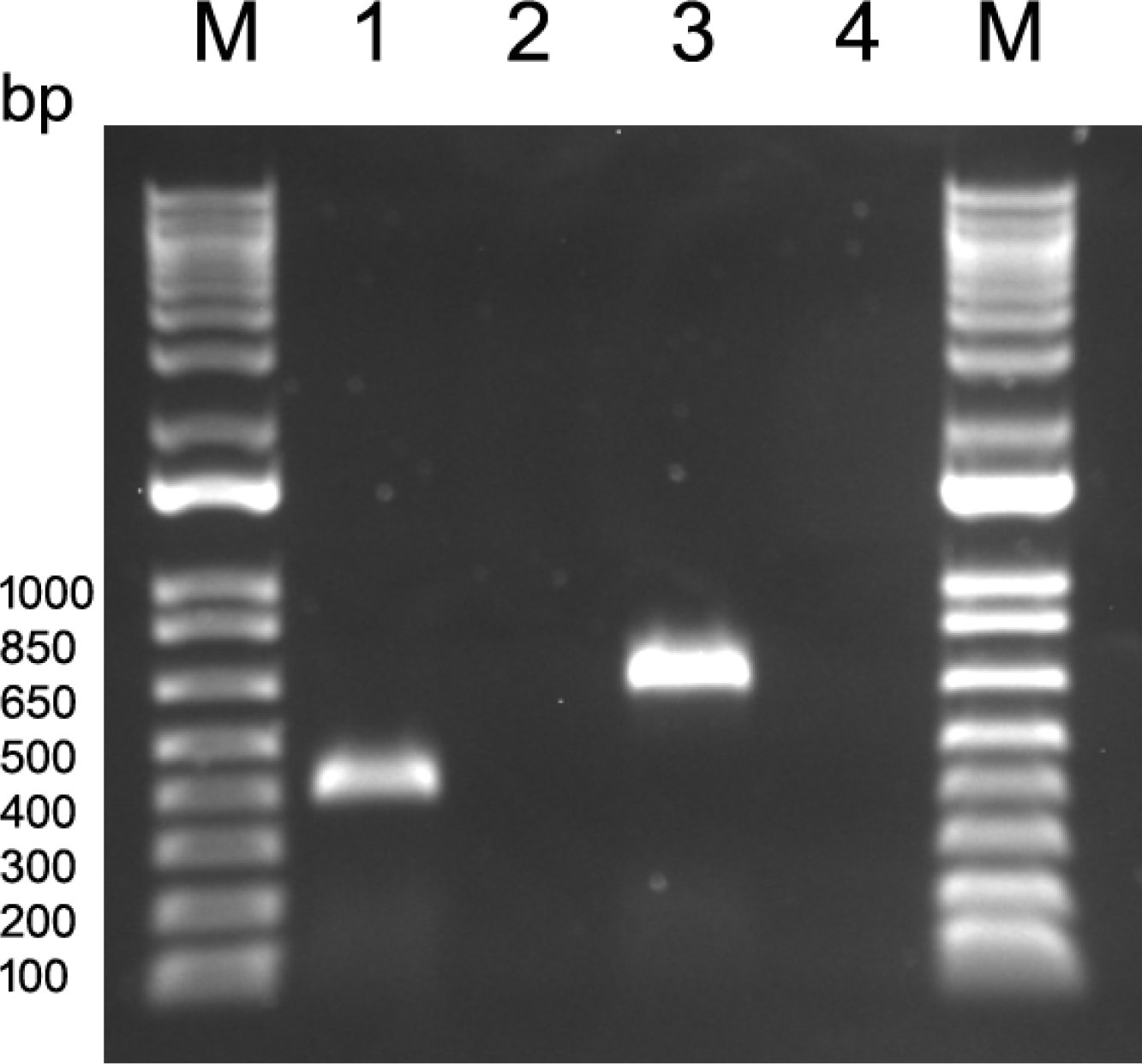
RT-PCR assay with symptomatic leaves of mechanically inoculated alfalfa plants. **M**, 1 Kb Plus DNA Ladder (Thermo Fisher Scientific, Waltham, MA USA). **1**, PCR amplicon obtained from the symptomatic leaves of alfalfa cv. CUF101 using primer pair LN1077/LN1078. **2**, no PCR amplicon was obtained from control reaction using the same primers. **3**, PCR amplicon obtained from the symptomatic alfalfa leaves using primer pair LN1079/LN1079. **4**, no PCR amplicon was obtained from control reaction with the same primers.

### Serological examinations

In Western blot (WB) assays, virus-infected tissues did not react with broad spectrum monoclonal antibodies (Abs) to potyviruses (PTY 1), [28], (results not shown). However, they appeared to weakly cross-react with polyclonal antisera to SPMMV (DSMZ, Braunschweig, Germany) when an indirect WB assay was used (membranes were probed with primary Abs to SPMMV followed by secondary Ab-alkaline phosphatase conjugate) was used. This reaction was accompanied by a high background and non-specific reactivity, common for indirect serological detection protocols (**Fig. S1A**). When membranes were probed with SPMMV primary Abs conjugated with AP (direct assay), the only reactive band was derived from the purified viral preparation, likely due to the large concentration of the viral coat protein rather than to cross-reactivity with SPMMV antiserum. No cross-reactivity was observed with virus-infected plant tissues (**Fig S1B**). This experiment indicated a possible distant serological relationship between the novel virus and a type species of the genus *Ipomovirus*. Although the percent amino acid identity between the SPMMV CP and presumptive CP of the alfalfa ipomovirus is quite low (31%) and is far below the identity necessary for classification into the genus *Ipomovirus*, the latter may possess some epitopes recognized by the SPMMV polyclonal Abs.

## Discussion

We report here the identification and characterization of a new virus infecting alfalfa (*Medicago sativa* L.) and belonging to the family *Potyviridae*. Genomic fragments of the same virus were previously found in mature alfalfa seeds [15]. Since they shared some homology (∼30%) to several potyvirids from different genera of the family, the virus was provisionally named alfalfa-associated potyvirus 1 (AaPV1) [15].

In this study, based on the complete genome sequence and structure, phylogenetic relationships, TEM observations, symptomatology on its primary agricultural host, and in part by serological cross-reactivity, we propose a more descriptive name for the virus, alfalfa vein mottling virus (AVMV). Additionally, our results indicate that AVMV should be reclassified as a novel species related to but distinct from viruses of the genus *Ipomovirus*. Furthermore, distinct clustering of AVMV in the phylogenetic tree and low identity levels with other members of the genus *Ipomovirus* suggest that alfalfa vein mottling virus likely represents a new species of a unique taxon most closely related to the genus *Ipomovirus*. We propose this new genus be named Alvemovirus.

The characteristic symptoms caused by the virus on indicator plant *Nicotiana benthamiana* and on its agricultural host alfalfa, confirm pathogenicity of AVMV on these host species. Successful mechanical inoculation of *M. sativa* with virus purified from the indicator plant followed by RT-PCR detection of the AVMV in mechanically inoculated alfalfa, satisfies Koch’s postulates and establishes the virus as cause of the symptoms observed on *M. sativa* plants.

Viruses of the genus *Ipomovirus*, to which AVMV is presumably related but distinct, infect a wide range of hosts, including economically important crops such as tomato, cucumber, melon, squash, sweet potato, watermelon, and others. To our knowledge, ipomoviruses or closely related viruses have not previously been identified from alfalfa. Our findings indicate that *M. sativa* can be a natural reservoir for the AVMV and thus, considering its possible relationship to ipomoviruses, which are transmitted by sweetpotato whitefly (*Bemisia tabaci*), it is possible that AVMV is also whitefly-transmitted and could infect other crops growing in the vicinity. Therefore, further studies will examine incidence of AVMV in additional regional crop and/or weed species and the potential for AVMV transmission by *B. tabaci*. Since all alfalfa samples used in this study were collected within 100 m of the melon fields in an area with exceptionally high populations of the whitefly, *B. tabaci* MEAM1, (**Table 1**), further testing this melon would likely clarify the virus origin and host range.

## Funding information

This study was supported by the United States Department of Agriculture, Agricultural Research Service (CRIS number 8042-21000-300-00D) and partially by the National Plant Disease Recovery System (NPDRS) grants to LGN.

## Author contributions

Conceptualization: LGN. Sample collection and evaluation: JCP and WMW; Methodology: OAP, LGN, SG. Data analysis: SG and LGN. Experimentation: LGN and OAP. Writing – original draft preparation: LGN. Writing – review and editing: all authors.

## Data Availability Statement

The genomic sequence of the alfalfa vein mottling virus has been deposited in GenBank on 08/24/2023 under the accession number OR483877.

## Conflicts of interest

The authors declare that there are no conflicts of interest.

## Notes

### Competing Interest Statement

The authors have declared no competing interest.

### Summary of Updates

Discussion section updated. Minor grammatical and wording changes made.

