## Supplementary figures and images for "Alfalfa vein mottling virus, a novel potyvirid infecting *Medicago sativa* L"

### Fig.S1A, Fig.S1B

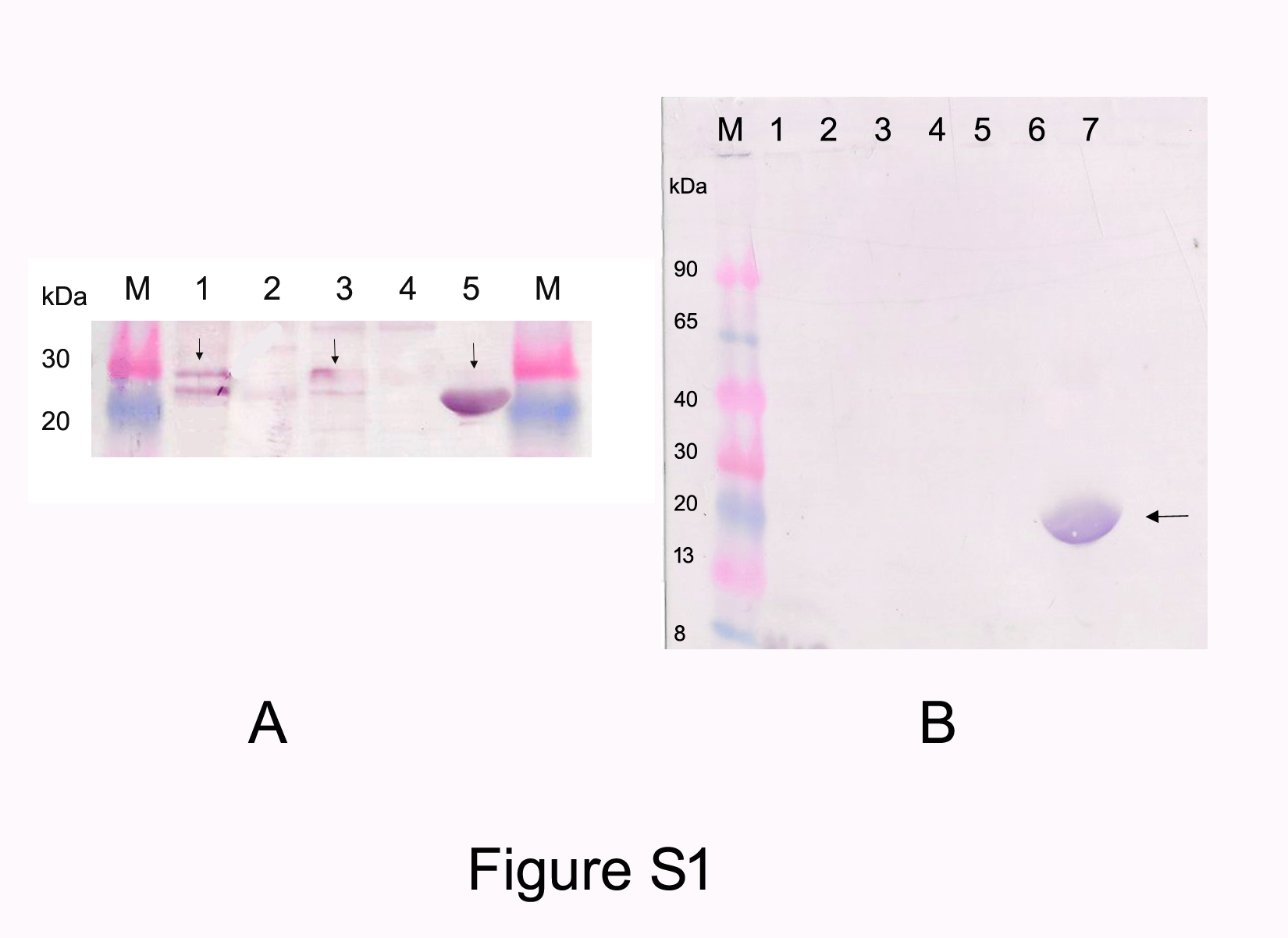
